# Pregnancy-Dependent Cardioprotection via GPER Activation in Dahl Salt-Sensitive Rats

**DOI:** 10.1101/2025.04.01.646468

**Authors:** Allan K. N. Alencar, Kenneth F. Swan, Mistina M. Manoharan, Christopher A. Natale, Sarah H. Lindsey, Gabriella C. Pridjian, Michael R. Garrett, Carolyn L. Bayer

## Abstract

**Background:** Preeclampsia is a hypertensive disorder of pregnancy that affects multiple organs, including the heart, increasing long-term cardiovascular risks for both the mother and offspring. While the G protein-coupled estrogen receptor (GPER) has cardioprotective effects, its role in pregnancy-associated cardiac dysfunction, particularly in chronic hypertension, remains unclear, given the significant physiological adaptations that occur during pregnancy, including hormonal fluctuations and hemodynamic changes. This study investigated whether LNS8801, a selective and orally bioavailable GPER agonist, could improve cardiac function in virgin and pregnant Dahl salt-sensitive (SS/Jr) rats, a model of chronic hypertension exacerbated by pregnancy.

**Methods:** Female Dahl SS/Jr rats, both virgin and pregnant, were randomized into four groups: Virgin + Vehicle, Virgin + LNS8801, Pregnant + Vehicle, and Pregnant + LNS8801. LNS8801 (800 µg/kg/day, given orally) was administered in pregnant rats from gestational day (GD) 9 to 20 and for an equivalent period in virgin controls. Cardiac function was assessed via echocardiography, including speckle-tracking strain analysis and conventional systolic and diastolic parameters. Mean arterial pressure and proteinuria were also measured.

**Results:** LNS8801 significantly improved cardiac function in pregnant Dahl SS/Jr rats, enhancing global longitudinal, circumferential, and radial strain, as well as increasing systolic function. Additionally, LNS8801 enhanced diastolic function, improving left ventricular compliance (E/A ratio) and early mitral annular velocity (e′), while reducing left ventricular filling pressures (E/e′ ratio). In contrast, LNS8801 had no significant effects on cardiac function and blood pressure in virgin Dahl SS/Jr rats, suggesting that pregnancy-related adaptations may enhance GPER-mediated cardioprotection. LNS8801 treatment significantly reduced proteinuria in both virgin and pregnant rats, indicating a pregnancy-independent renal protective effect.

**Conclusion:** This study highlights the importance of pregnancy-specific adaptations in shaping the cardiovascular effects of GPER activation. While LNS8801 demonstrated cardioprotective and antihypertensive benefits in pregnant Dahl SS/Jr rats, its effects were absent in virgin animals, underscoring the influence of the physiological and hormonal environment on GPER-mediated responses. These findings provide a foundation for further exploration of GPER as a therapeutic target for pregnancy-associated cardiovascular dysfunction and preeclampsia, reinforcing the need for pregnancy-specific approaches in drug development.

## Introduction

Preeclampsia is a hypertensive condition that occurs in 5%–8% of pregnancies, leading to complications that can affect various organs, including the heart, kidneys, brain, and liver^1,2^. This disorder poses significant health risks to both the mother and the fetus, potentially causing lasting damage to these vital organs if not properly managed^3,4^.

Preeclampsia primarily stems from abnormalities in placental development, where disruptions in normal vascular formation can result in placental insufficiency. This insufficiency then contributes to various maternal symptoms and clinical complications ^5,6^. Classically, when pregnant women are diagnosed with preeclampsia, they present with new-onset hypertension and proteinuria after 20 weeks of gestation^7^. Preeclampsia can also be classified as superimposed when it occurs in individuals with pre-existing chronic hypertension, where pregnancy exacerbates the condition and increases the risk of maternal and fetal complications^7,8^.

The primary treatment for preeclampsia focuses on stabilizing the mother and delivering the fetus and placenta early, often resulting in preterm birth and low birth weight, which heighten cardiovascular risks in offspring^9^. Developing therapies that address the full range of symptoms remains challenging, as current interventions are limited, with placental delivery being the only definitive solution^10^. Improved, lasting treatments are urgently needed. Chronic hypertension significantly raises the risk of developing preeclampsia^11^, and this combination can further strain the heart, leading to increased cardiac issues^12^. The rising prevalence of related comorbidities, such as obesity, exacerbates both hypertension and the risk of heart dysfunction, adding to the overall burden of maternal health^13^.

Selective activation of the G protein-coupled estrogen receptor (GPER) has been extensively studied in models of chronic hypertension, highlighting its potential as a therapeutic target^14–17^. For example, GPER stimulation with the agonist G-1 reduces blood pressure in estrogen-depleted chronic hypertensive mRen2.Lewis rats^14^.

Additionally, G-1 improves myocardial relaxation and reduces cardiac myocyte hypertrophy in female mRen2.Lewis rats on a high-salt diet^15^. However, these studies were conducted in ovariectomized animals, where the absence of endogenous estrogen minimizes competition for G-1 binding to GPER. In non-pregnant, non-ovariectomized hypertensive rats, G-1’s effects are attenuated or absent, likely due to pharmacodynamic competition between endogenous estrogen and exogenous G-1^14–17^. In non-ovariectomized Dahl SS/Jr rats on a high-salt diet, G-1 ameliorated kidney injury and reduced proteinuria without significantly altering blood pressure^17^.

Previous research has demonstrated that Dahl SS/Jr rats naturally develop a preeclampsia-like condition during pregnancy, characterized by elevated blood pressure, worsening proteinuria, fetal growth restriction, and subclinical cardiac dysfunction, superimposed on pre-existing chronic hypertension^18,19^. While we recently demonstrated that GPER activation with G-1 improves cardiac function in reduced uterine perfusion pressure dams^20^, a model of placental ischemia-associated hypertension, whether pregnancy itself modifies GPER’s cardioprotective effects remains unclear.

Pregnancy induces significant hemodynamic adaptations, including increased cardiac output, blood volume, and vascular remodeling, which could influence the response to GPER activation. Additionally, the presence of high endogenous estrogen levels during pregnancy^21^ may alter the pharmacodynamic effects of GPER agonists compared to the non-pregnant state. To address this gap, we investigated the effects of LNS8801, a selective, enantiomerically pure, and orally bioavailable GPER agonist that is currently undergoing clinical trials for cancer^22^ on cardiac function in Dahl SS/Jr rats, comparing virgin and pregnant animals. Investigating LNS8801 in both pregnant and non-pregnant hypertensive rats is necessary for understanding whether its effects are influenced by the unique hormonal and hemodynamic environment of pregnancy or if they persist independently of reproductive status, providing new insights into the potential of LNS8801 as a therapeutic target for pregnancy-associated cardiac dysfunction.

## Material and Methods

### Data Availability

Further methodological details are available in the Supplemental Material. Data supporting the reported studies are available from the corresponding author upon reasonable request.

### Animals

Virgin female Dahl SS/Jr rats (ages ranging between 18 and 22 weeks) were obtained from a colony maintained at the University of Mississippi Medical Center. All rats were fed a low-salt rodent chow (TD7034, 0.3% NaCl; Harlan Teklad, Madison, WI) and housed in conventional housing with two to four rats per cage in a controlled environment with a temperature of 21°C, maintained on a 12-hour light/dark cycle, with food and tap water provided ad libitum. For the pregnant groups, timed breeding was performed using breeding screens and the presence of a vaginal plug was indicative of gestational day (GD) 1. The Institutional Animal Care and Use Committee at Tulane University approved all animal studies, which were conducted in accordance with the guidelines outlined in the NIH Guide for the Care and Use of Laboratory Animals.

### Experimental treatment protocol

In this study, rats were randomly assigned to four experimental groups: Virgin + Vehicle (n = 5), Virgin + LNS8801 (Linnaeus Therapeutics; n = 6), Pregnant + Vehicle (n = 7), and Pregnant + LNS8801 (n = 6). Previous studies have demonstrated that the racemic GPER agonist G-1 significantly reduces blood pressure^14^ and confers cardioprotective effects^23^ when administered subcutaneously at 400 µg/kg/day in female rats. However, in a pilot study conducted in our laboratory, oral administration of LNS8801, the purified active enantiomer of G-1, at the same dose (400 µg/kg/day) in Dahl SS/Jr rats exhibiting superimposed preeclampsia, significantly reduced blood pressure but did not yield significant cardiac improvements compared to vehicle-treated controls. Therefore, a higher oral dose of 800 µg/kg/day was selected for the current work. LNS8801 was initially dissolved in dimethyl sulfoxide at 10 µg/µL and then 24 µL (240 µg) was mixed with 10% ethanol and 80% soybean oil (Spectrum Chemical, Gardena, USA). A total of 200 µL of this solution was administered via oral gavage, delivering a target dose of 800 µg/kg/day. The control groups received 200 µL of the vehicle, which was the same solution used to dissolve LNS8801. LNS8801 or the vehicle were administered for 12 days, from GD9 to GD20 in pregnant rats, and for an equivalent period in age-matched virgin controls. Rats were weighed daily, and the dosages of LNS8801 were adjusted appropriately.

### Echocardiography

On GD18, or the corresponding day in age-matched virgin animals, rats were positioned supine on a heated pad to maintain a body temperature of 37°C, with anesthesia induced by 3% isoflurane/oxygen mixture and reduced to 1-2% during the imaging procedure. Depilatory cream was applied to the ventral thorax for clear imaging, and electrocardiogram monitoring was performed using limb electrodes. Pre-warmed ultrasound gel was applied to avoid cold stress. Imaging was performed using the Vevo 2100 Imaging System (Visual Sonics, Toronto, Canada) with a 20 MHz ultra-high-frequency linear-array transducer. Parasternal long-axis images were first obtained, followed by short-axis images at the mid-papillary level. Speckle-tracking technology was utilized to assess global longitudinal strain in the long-axis, circumferential strain in the short-axis, and radial strain in both long- and short-axis to detect subclinical cardiac dysfunction. B-mode videos with a frame rate exceeding 200 fps were selected for strain analysis. For consistency, three consecutive cardiac cycles between respiratory cycles were analyzed, with endocardial and epicardial borders traced semi-automatically using multiple tracking points. Speckle-tracking strain analyses were performed offline using VevoStrain software (Visual Sonics, Toronto, Canada), following established protocols^24^. Parasternal long- and short-axis B-mode views were used to evaluate conventional left ventricular (LV) systolic parameters, including ejection fraction, fractional shortening, and cardiac output, which was calculated using Simpson’s method. LV structural measurements, such as wall thickness and diameter, were obtained via M-mode at the mid-papillary level, with averages taken over five cardiac cycles. Mitral inflow measurements of early and late filling velocities (E and A waves, respectively) were obtained using pulsed Doppler, with the sample volume placed at the tips of mitral leaflets from an apical four-chamber orientation. Tissue Doppler imaging to assess mitral valve septal annular velocities (e′ and a′ waves) was also obtained from the four-chamber view. All measured and calculated systolic and diastolic indices are represented as the average of at least five consecutive cardiac cycles to minimize beat-to-beat variability. To assess both intra- and interobserver variability, we randomly selected 20 images. These images were blindly analyzed twice at 1-month intervals by one trained observer and independently analyzed by a second trained observer.

### Urinary Protein Excretion

The rats were placed in metabolic cages for 24-hour urine collection at the end of pregnancy (from GD18 to 19) or on the corresponding days in age-matched virgin animals. Urinary protein concentration was measured using the Bradford Assay (BioRad Laboratories, Hercules, CA, USA), and proteinuria was defined as >20 mg/24 h.

### Terminal Mean Arterial Pressure Measured under Anesthesia

On GD20, at the time of tissue harvesting, rats were anesthetized with isoflurane, and a saline-filled catheter was surgically placed in the left common carotid artery to measure mean arterial pressure, following established procedures^25^. For virgin animals, the procedure was performed on the corresponding day to match the experimental timeline. After catheter insertion, a heparinized saline flush was used to maintain patency.

Anesthetized animals were positioned on a heated physiological platform, and the catheter was connected to a pressure transducer linked to a data acquisition system (PowerLab, ADInstruments, Colorado Springs, CO) for real-time blood pressure recordings. All rats were kept under light, stable anesthesia with controlled respiratory and heart rates by adjusting the isoflurane concentration. After the experiments, animals were euthanized by decapitation under isoflurane anesthesia.

### Tissue collection

On gestational GD20, tissue collection was carried out. The number of viable and resorbed fetuses in each uterine horn was recorded. For virgin animals, tissue collection took place on the equivalent experimental day. Placental and fetal weights for each viable fetus were measured, and the averages were calculated per litter. The fetal-to-placental weight ratio was determined for each unit before being averaged per litter. Hearts were carefully blotted dry on filter paper and then weighed.

### Statistics

Sample sizes were determined from an a priori power analysis performed in G*Power Software (Heinrich-Heine-Universität, Dusseldorf, Germany). All data are shown as means ± SEM, with “n” representing the number of individual experiments. Statistical analyses were performed on GraphPad Prism 10.4.1 (GraphPad Software, San Diego, CA). The Shapiro-Wilk normality test was used to assess the Gaussian distribution (α = 0.05). Two-way ANOVA was used to assess the main effects of treatment and pregnancy, as well as their interaction, followed by Tukey’s multiple comparisons test. For fetal and placental outcomes, which involved comparisons within pregnant animals only, unpaired two-tailed t-tests were applied. Pearson’s correlation analysis (r) was used to evaluate associations between two variables. A *p*-value < 0.05 was considered statistically significant.

## Results

### Oral administration of LNS8801 improves cardiac strain parameters in pregnant Dahl SS/Jr rats but not in virgins

Cardiac strain reflects myocardial deformation during the cardiac cycle. More negative longitudinal and circumferential strain values indicate greater myocardial shortening, while more positive radial strain values indicate greater myocardial wall thickening. In both cases, greater deformation (% change) is associated with improved cardiac function. Our laboratory previously demonstrated that LV global longitudinal strain in healthy pregnant Sprague-Dawley rats is -20.9 ± 0.6% during late pregnancy^26^.

Treatment with LNS8801 significantly restored LV longitudinal strain in long axis on GD18 in pregnant Dahl SS/Jr rats (-21.8 ± 1.2%) compared to vehicle-treated pregnant controls (-11.6 ± 1.5%; Figure 1A), indicating enhanced myocardial deformation.

**Figure 1:**
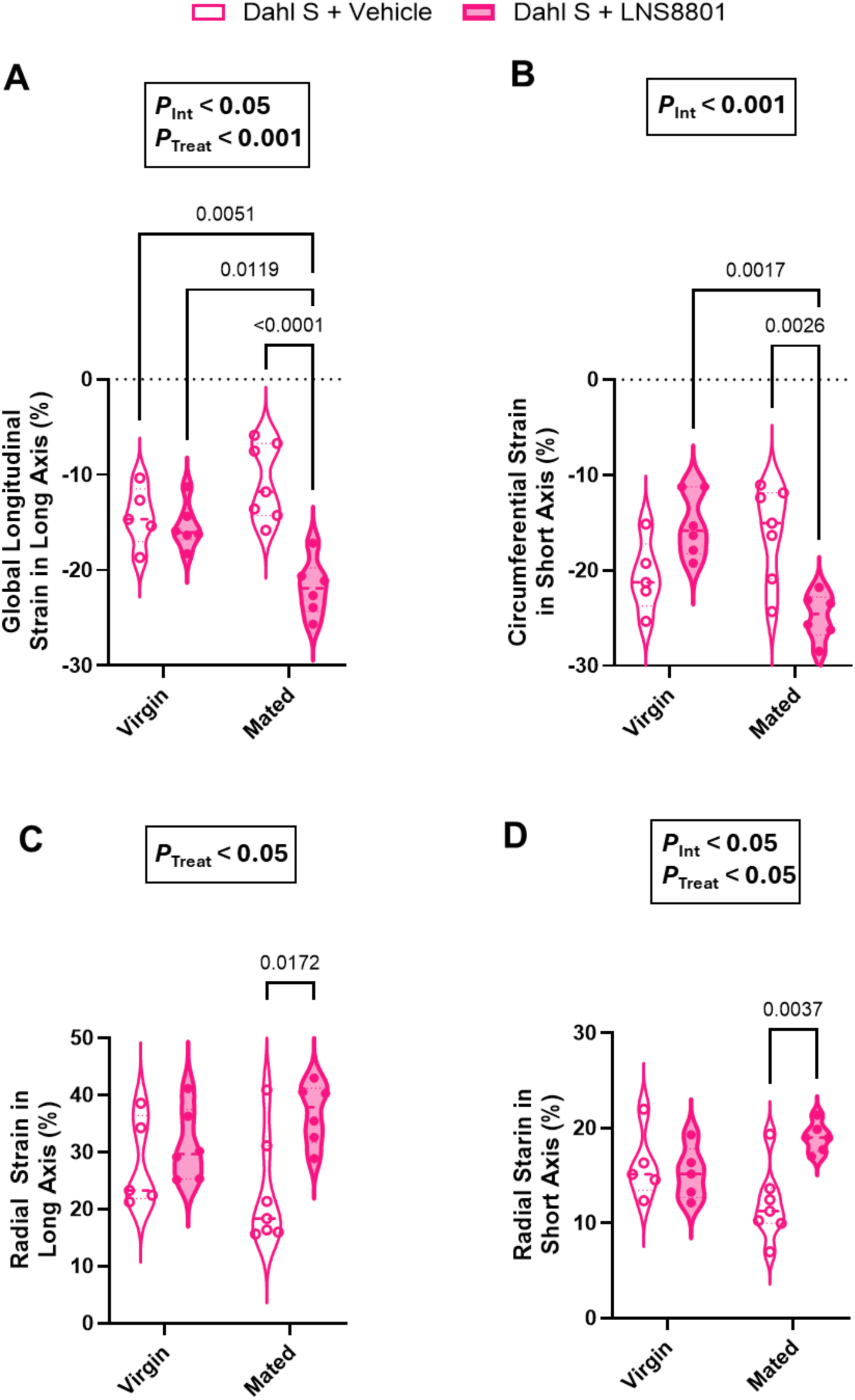
LNS8801 administration differentially affects myocardial strain in pregnant and virgin Dahl SS/Jr rats. (**A**) Global longitudinal strain in the long axis, (**B**) circumferential strain in the short axis, (**C**) radial strain in the long axis, and (**D**) radial strain in the short axis were assessed on gestational day 18. LNS8801 treatment significantly improved all strain parameters in pregnant rats compared with vehicle-treated controls, indicating reduced myocardial deformation. In contrast, no significant differences were observed between LNS8801- and vehicle-treated virgin rats, suggesting that pregnancy-specific adaptations may enhance GPER-mediated cardioprotection. Significant main effects of treatment and/or interaction (pregnancy × treatment) from two-way ANOVA are shown above each graph, and Tukey’s post hoc comparisons are indicated between groups. Data are presented as mean ± SEM. Individual points represent biological replicates (n = 5 for Virgin + Vehicle, n = 6 for Virgin + LNS8801, n = 7 for Mated + Vehicle, and n = 6 for Mated + LNS8801).

Additionally, global longitudinal strain in LNS8801-treated pregnant rats was significantly lower than in both virgin rats treated with vehicle and those treated with LNS8801 (Figure 1A). No significant differences in global longitudinal strain were observed between the virgin groups. These results were supported by two-way ANOVA, which showed a significant main effect of treatment (*F*(1, 20) = 20.88, *p* = 0.0002, 35.48% of variation) and a significant interaction between pregnancy and treatment (*F*(1, 20) = 14.24, *p* = 0.0012, 24.18%), but no main effect of pregnancy alone (*F*(1, 20) = 1.214, *p* = 0.2835, 2.06%; Figure 1A). We additionally measured LV circumferential strain in the short axis (Figure 1B). In pregnant Dahl SS/Jr rats, treatment with LNS8801 resulted in significantly more negative strain values compared to vehicle-treated pregnant controls, indicating enhanced myocardial deformation. No significant differences in circumferential strain were observed between virgin groups, nor between virgin and LNS8801-treated pregnant rats (Figure 1B). Two-way ANOVA for circumferential strain revealed a significant interaction between pregnancy and treatment (*F*(1, 20) = 20.49, *p* = 0.0002, 45.59% of variation), but no significant main effects of pregnancy (*F*(1, 20) = 2.457, *p* = 0.1327, 5.47%) or treatment (*F*(1, 20) = 1.169, *p* = 0.2924, 2.60%) were observed (Figure 1B). LV radial strain in both the long and short axes was significantly greater in the hearts of Dahl SS/Jr rats with superimposed preeclampsia treated with LNS8801, compared to vehicle-treated pregnant controls (Figures 1C and 1D, respectively). No significant differences were detected between virgin groups, nor between virgin and LNS8801-treated pregnant rats. Two-way ANOVA for radial strain in the long axis revealed a significant main effect of treatment (*F*(1, 20) = 7.573, *p* = 0.0123, 24.10% of variation), but no significant effects of pregnancy (*F*(1, 20) = 0.0053, *p* = 0.9426, 0.017%) or interaction (*F*(1, 20) = 2.948, *p* = 0.1015, 9.38%; Figure 1C). For radial strain in the short axis, two-way ANOVA showed a significant main effect of treatment (*F*(1, 19) = 5.491, *p* = 0.0302, 15.46% of variation) and a significant interaction between treatment and pregnancy (*F*(1, 19) = 8.868, *p* = 0.0077, 24.97%), while the effect of pregnancy alone was not significant (*F*(1, 19) = 0.0164, *p* = 0.8994, 0.046%; Figure 1D). Circulating levels of cardiac troponin I were measured and are presented in Figure S1A. Troponin I levels were significantly lower in pregnant rats treated with LNS8801 (0.2 ± 0.03 ng/mL) compared to virgin rats treated with LNS8801 (0.8 ± 0.02 ng/mL; Figure S1A). No significant differences were observed between any other groups (Figure S1A). Two-way ANOVA revealed a significant interaction between pregnancy and LNS8801 treatment (*F*(1, 17) = 9.579, *p* = 0.0066), as well as a main effect of pregnancy (*F*(1, 17) = 11.28, *p* = 0.0037). These factors accounted for 24.21% and 28.51% of the total variation in circulating cardiac troponin I levels, respectively. No significant main effect of treatment alone was observed (*F*(1, 17) = 0.0674, *p* = 0.7983), contributing minimally to the variation (0.17%). Additionally, circulating cardiac troponin I levels showed a significant correlation with global longitudinal strain (Figure S1B). Together, these findings support the concept that pregnancy modifies the cardiac response to GPER stimulation.

### Diastolic function is improved by GPER activation during pregnancy but remains unchanged in virgin Dahl SS/Jr rats

Transmitral pulsed-wave and myocardial tissue Doppler indices of diastolic function are shown in Figure 2. Visual representations of the transmitral (E wave and A wave) and myocardial tissue Doppler (e’ and a’) waves are depicted in Figure 2A. Early transmitral filling velocities (E wave) in healthy pregnant Sprague-Dawley rats are 806.7 ± 19.1 mm/s^26^. In this study, E wave velocities (Table 1) significantly increased from 632.2 ± 52.0 mm/s in pregnant Dahl SS/Jr rats treated with the vehicle, from 429.8 ± 10.4 mm/s in virgin rats treated with the vehicle, and from 512.0 ± 14.5 mm/s in virgin animals treated with LNS8801 to 863.2 ± 55.3 mm/s in pregnant animals that received 12 days of oral administration of LNS8801 (Table 1). Additionally, E wave velocities were significantly higher in pregnant vehicle-treated rats compared to virgin vehicle-treated controls (Table 1). No significant difference was observed between virgin animals treated with vehicle and those treated with LNS8801. Two-way ANOVA for E wave velocities revealed a significant effect of pregnancy (*F*(1, 20) = 42.35, *p* < 0.0001), accounting for 55.23% of the total variation, and a significant effect of treatment (*F*(1, 20) = 13.55, *p* = 0.0015, 17.68%). However, the interaction between pregnancy and treatment was not statistically significant (*F*(1, 20) = 3.059, *p* = 0.0956), contributing to 3.99% of the variation. The ratio of early-to-late LV filling velocities (E/A ratio) was significantly higher in pregnant Dahl SS/Jr rats treated with LNS8801 (1.61 ± 0.1) compared to pregnant vehicle-treated rats (1.05 ± 0.1), virgin vehicle-treated rats (0.85 ± 0.06), and virgin rats treated with LNS8801 (0.81 ± 0.04; Figure 2B). No significant differences in E/A ratio were observed between the virgin groups (Figure 2B). Two-way ANOVA revealed significant main effects of pregnancy (*F*(1, 20) = 32.58, *p* < 0.0001, 45.13% of variation) and treatment (*F*(1, 20) = 8.763, *p* = 0.0077, 12.14%), along with a significant interaction between pregnancy and treatment (*F*(1, 20) = 11.31, *p* = 0.0031, 15.67%; Figure 2B). These results suggest that GPER activation improves LV compliance altered by superimposed preeclampsia. Additionally, the tissue Doppler measure of myocardial relaxation, mitral annular descent velocity (e′), was significantly increased in LNS8801-treated pregnant rats (44.5 ± 4.3 mm/s) compared to their pregnant controls (30.6 ± 2.0 mm/s; Table 1), as well as to virgin rats treated with vehicle (28.6 ± 2.3 mm/s) or LNS8801 (30.6 ± 1.5 mm/s; Table 1). No significant difference was observed between the virgin groups (Table 1). This was confirmed by two-way ANOVA, which showed significant main effects of pregnancy (*F*(1, 20) = 9.135, *p* = 0.0067, 20.96%) and treatment (*F*(1, 20) = 9.251, *p* = 0.0064, 21.22%), as well as a significant interaction (*F*(1, 20) = 5.247, *p* = 0.0330, 12.04%), indicating that LNS8801 improves myocardial relaxation only in the context of cardiac dysfunction worsened by superimposed preeclampsia. The echo-derived LV filling pressures, represented by the ratio of early filling to mitral annular descent velocities (E/e′ ratio), were significantly lower in pregnant rats treated with LNS8801 (18.4 ± 1.0) compared to pregnant rats treated with vehicle (23.0 ± 1.6; Figure 2C). Additionally, vehicle-treated pregnant rats exhibited higher E/e′ ratios than both virgin rats treated with vehicle (15.4 ± 1.3) and those treated with LNS8801 (16.8 ± 0.8; Figure 2C). No significant difference was observed between the two virgin groups (Figure 2C). Two-way ANOVA identified a significant interaction between pregnancy and treatment (*F*(1, 20) = 5.961, *p* = 0.0240, 14.21%) and a main effect of pregnancy (*F*(1, 20) = 12.56, *p* = 0.0020, 29.95%), while the main effect of treatment was not significant (*F*(1, 20) = 1.958, *p* = 0.1771, 4.67%).

**Figure 2:**
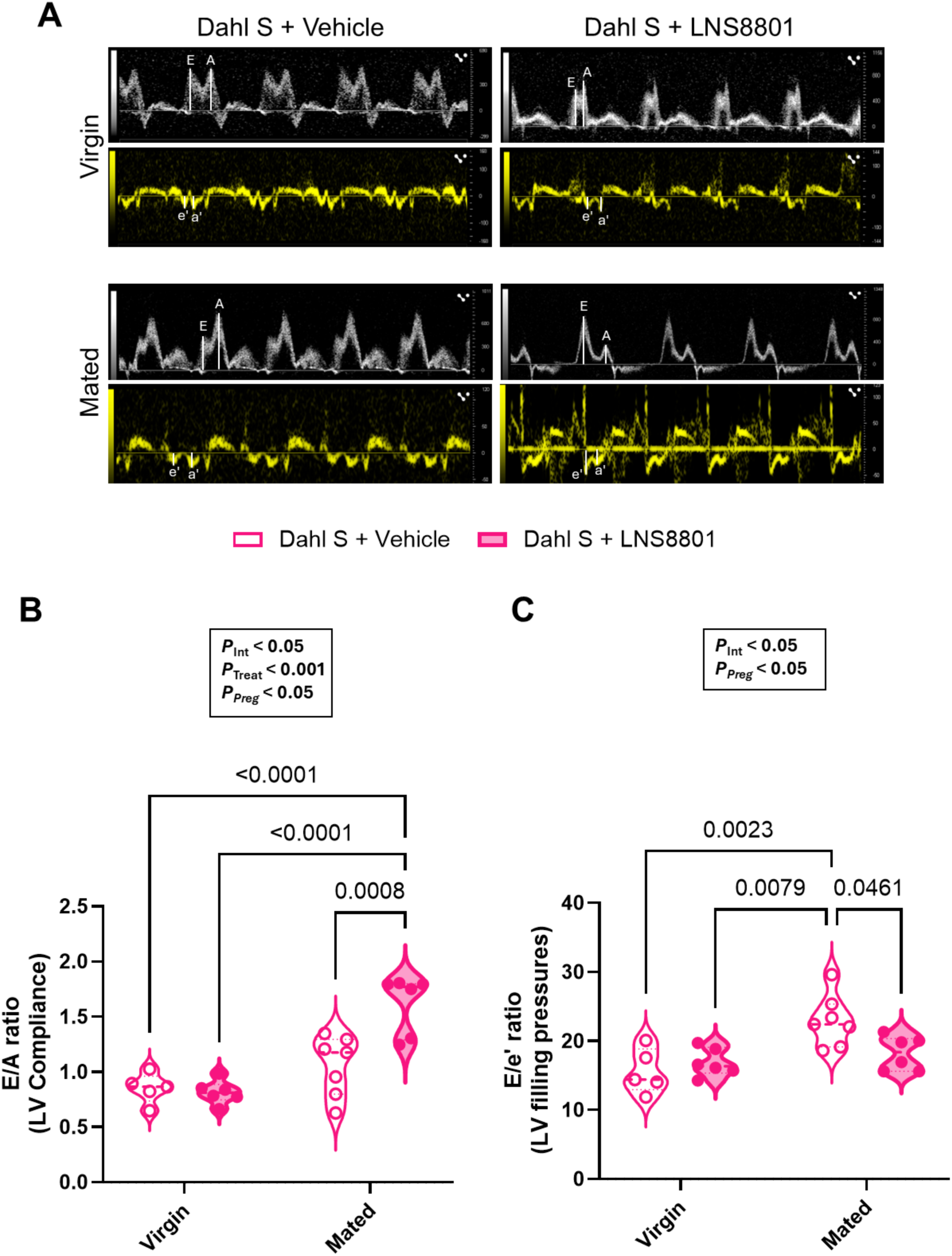
LNS8801 improves diastolic function in pregnant but not virgin Dahl SS/Jr rats. (**A**) Representative images of pulse wave Doppler of the mitral valve in four-chamber view, showing early (E) and late (A) filling waves and early (e’) and late (a’) diastolic mitral annular velocities, which were measured using tissue Doppler imaging. (**B**) The E/A ratio was significantly increased in pregnant rats treated with LNS8801 compared to all other groups, including pregnant vehicle-treated rats, virgin vehicle-treated rats, and virgin rats treated with LNS8801. No differences were observed between the virgin groups. (**C**) The E/e′ ratio was significantly lower in pregnant rats treated with LNS8801 compared to pregnant vehicle-treated controls. Pregnant vehicle-treated rats exhibited higher E/e′ ratios than both virgin groups, whereas no differences were detected between virgins treated with vehicle or LNS8801.Significant main effects of treatment and/or interaction (pregnancy × treatment) from two-way ANOVA are shown above each graph, and Tukey’s post hoc comparisons are indicated between groups, with significance set at (*p* < 0.05). Data are presented as mean ± SEM. Individual points represent biological replicates (n = 5 for Virgin + Vehicle, n = 6 for Virgin + LNS8801, n = 7 for Mated + Vehicle, and n = 6 for Mated + LNS8801).

**Table 1:**
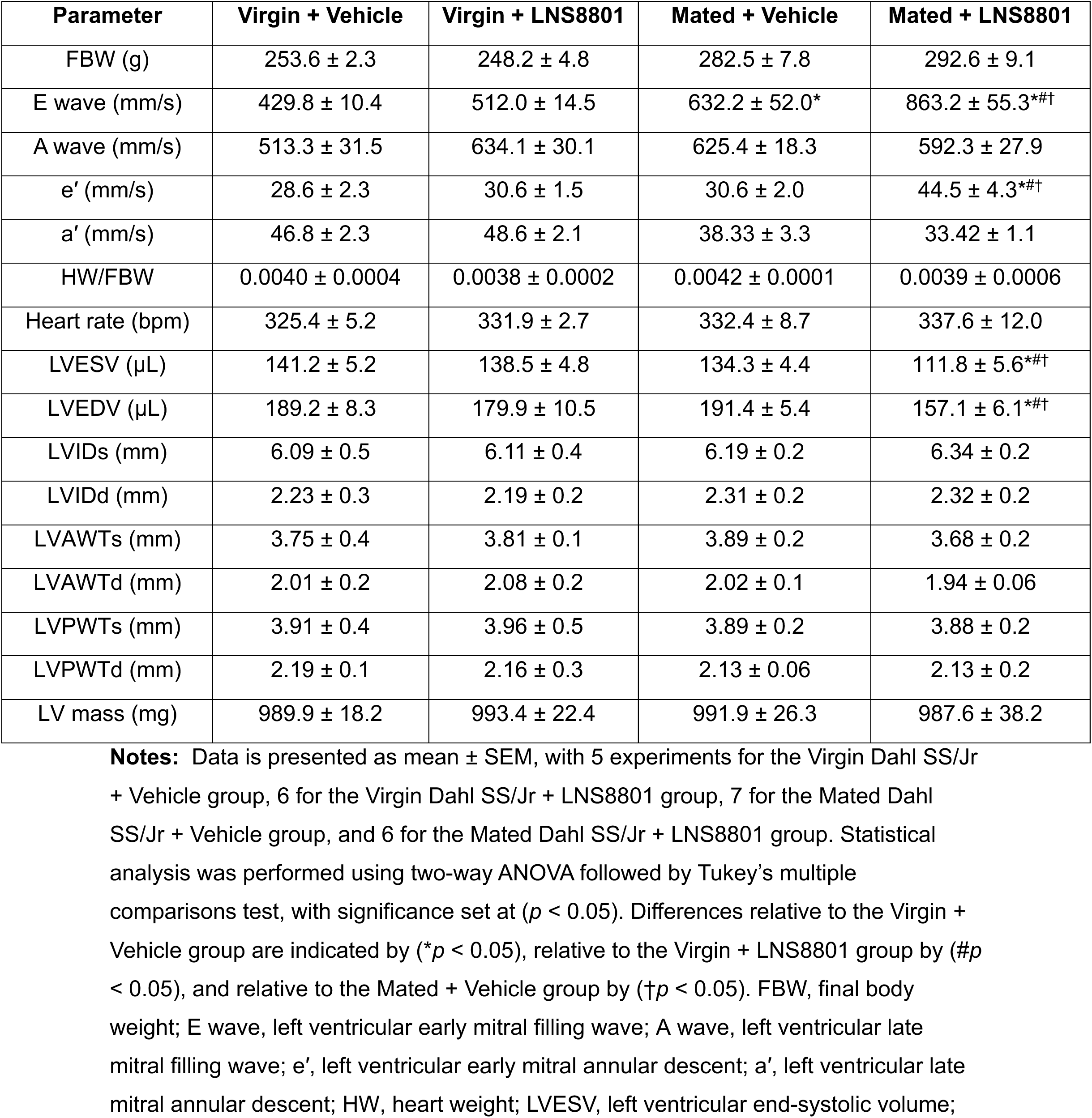
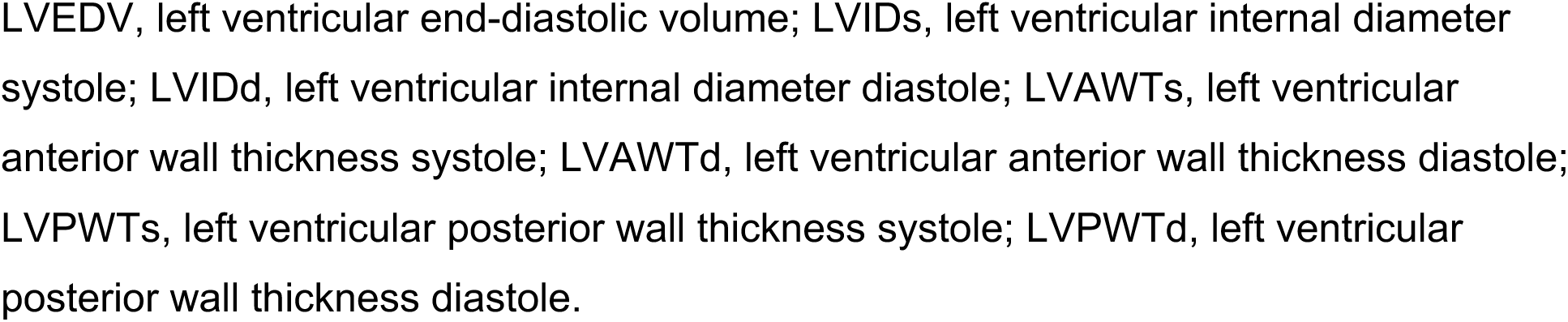
Echocardiographic and morphometric parameters in Dahl SS/Jr rats treated with LNS8801.

| Parameter | Virgin + Vehicle | Virgin + LNS8801 | Mated + Vehicle | Mated + LNS8801 |
| --- | --- | --- | --- | --- |
| FBW (g) | 253.6 ± 2.3 | 248.2 ± 4.8 | 282.5 ± 7.8 | 292.6 ± 9.1 |
| E wave (mm/s) | 429.8 ± 10.4 | 512.0 ± 14.5 | 632.2 ± 52.0* | 863.2 ± 55.3*#† |
| A wave (mm/s) | 513.3 ± 31.5 | 634.1 ± 30.1 | 625.4 ± 18.3 | 592.3 ± 27.9 |
| e' (mm/s) | 28.6 ± 2.3 | 30.6 ± 1.5 | 30.6 ± 2.0 | 44.5 ± 4.3*#† |
| a' (mm/s) | 46.8 ± 2.3 | 48.6 ± 2.1 | 38.33 ± 3.3 | 33.42 ± 1.1 |
| HW/FBW | 0.0040 ± 0.0004 | 0.0038 ± 0.0002 | 0.0042 ± 0.0001 | 0.0039 ± 0.0006 |
| Heart rate (bpm) | 325.4 ± 5.2 | 331.9 ± 2.7 | 332.4 ± 8.7 | 337.6 ± 12.0 |
| LVESV (μL) | 141.2 ± 5.2 | 138.5 ± 4.8 | 134.3 ± 4.4 | 111.8 ± 5.6*#† |
| LVEDV (μL) | 189.2 ± 8.3 | 179.9 ± 10.5 | 191.4 ± 5.4 | 157.1 ± 6.1*#† |
| LVIDs (mm) | 6.09 ± 0.5 | 6.11 ± 0.4 | 6.19 ± 0.2 | 6.34 ± 0.2 |
| LVIDd (mm) | 2.23 ± 0.3 | 2.19 ± 0.2 | 2.31 ± 0.2 | 2.32 ± 0.2 |
| LVAWTs (mm) | 3.75 ± 0.4 | 3.81 ± 0.1 | 3.89 ± 0.2 | 3.68 ± 0.2 |
| LVAWTd (mm) | 2.01 ± 0.2 | 2.08 ± 0.2 | 2.02 ± 0.1 | 1.94 ± 0.06 |
| LVPWTs (mm) | 3.91 ± 0.4 | 3.96 ± 0.5 | 3.89 ± 0.2 | 3.88 ± 0.2 |
| LVPWTd (mm) | 2.19 ± 0.1 | 2.16 ± 0.3 | 2.13 ± 0.06 | 2.13 ± 0.2 |
| LV mass (mg) | 989.9 ± 18.2 | 993.4 ± 22.4 | 991.9 ± 26.3 | 987.6 ± 38.2 |

**Notes:** Data is presented as mean ± SEM, with 5 experiments for the Virgin Dahl SS/Jr + Vehicle group, 6 for the Virgin Dahl SS/Jr + LNS8801 group, 7 for the Mated Dahl SS/Jr + Vehicle group, and 6 for the Mated Dahl SS/Jr + LNS8801 group. Statistical analysis was performed using two-way ANOVA followed by Tukey's multiple comparisons test, with significance set at ( $p < 0.05$ ). Differences relative to the Virgin + Vehicle group are indicated by ( $*p < 0.05$ ), relative to the Virgin + LNS8801 group by ( $#p$ $< 0.05$ ), and relative to the Mated + Vehicle group by ( $†p < 0.05$ ). FBW, final body weight; E wave, left ventricular early mitral filling wave; A wave, left ventricular late mitral filling wave; e', left ventricular early mitral annular descent; a', left ventricular late mitral annular descent; HW, heart weight; LVESV, left ventricular end-systolic volume;
LVEDV, left ventricular end-diastolic volume; LVIDs, left ventricular internal diameter systole; LVIDd, left ventricular internal diameter diastole; LVAWTs, left ventricular anterior wall thickness systole; LVAWTd, left ventricular anterior wall thickness diastole; LVPWTs, left ventricular posterior wall thickness systole; LVPWTd, left ventricular posterior wall thickness diastole.

The significant interaction effect underscores the pregnancy-dependent nature of GPER-mediated diastolic improvement in this model.

### Effects of LNS8801 on systolic function and global cardiac performance in virgin and pregnant Dahl SS/Jr rats

LV systolic performance was quantified by measuring ejection fraction in virgin and pregnant Dahl SS/Jr rats (Figure 3A). In pregnant rats, ejection fraction was significantly higher in LNS8801-treated animals (91.0 ± 1.5%) compared to vehicle-treated controls (73.1 ± 3.5%), virgin rats treated with vehicle (78.5 ± 2.1%), and virgin rats treated with LNS8801 (80.7 ± 1.5%), whereas no significant differences were observed between virgin animals (Figure 3A). Two-way ANOVA confirmed a significant main effect of pregnancy (*F*(1, 20) = 10.77, *p* = 0.0037, 28.50%) and treatment (*F*(1, 20) = 5.597, *p* = 0.0282, 14.82%) on ejection fraction, with no significant interaction (*F*(1, 20) = 1.930, *p* = 0.1800, 5.10%). In pregnant Dahl SS/Jr rats treated with LNS8801, the LV stroke volume was significantly higher (209.9 ± 9.5 µL) compared to pregnant vehicle-treated controls (165.3 ± 15.3 µL), virgin vehicle-treated rats (158.3 ± 3.9 µL), and virgin rats treated with LNS8801 (164.5 ± 4.5 µL; Figure 3B). No significant differences were observed between the virgin groups (Figure 3B). Two-way ANOVA revealed significant main effects of pregnancy (*F*(1, 20) = 5.970, *p* = 0.0240, 17.19%) and treatment (*F*(1, 20) = 5.604, *p* = 0.0281, 16.13%) on LV stroke volume (Figure 3B), while the interaction between the two factors was not significant (*F*(1, 20) = 3.214, *p* = 0.0881, 9.25%). This increase in stroke volume was accompanied by a significant reduction in the LV end-systolic and end-diastolic volumes in pregnant rats treated with LNS8801, compared to pregnant vehicle-treated controls, as well as virgin rats treated with either vehicle or LNS8801 (Table 1). Two-way ANOVA confirmed significant main effects of pregnancy (*F*(1, 20) = 12.56, *p* = 0.0020, 27.84%) and treatment (*F*(1, 20) = 7.887, *p* = 0.0109, 17.47%) on LV end-systolic volume, as well as a significant interaction (*F*(1, 20) = 4.959, *p* = 0.0376, 10.99%). For LV end-diastolic volume, there were also significant main effects of pregnancy (*F*(1, 20) = 5.778, *p* = 0.0260, 9.52%) and treatment (*F*(1, 20) = 26.31, *p* < 0.0001, 43.37%), with a significant interaction (*F*(1, 20) = 7.690, *p* = 0.0117, 12.68%). In virgin rats, end-systolic volume and end-diastolic volume remained similar between groups (Table 1). These findings reinforce that LNS8801 significantly modulates cardiac volumes in a pregnancy-dependent manner. As a result, cardiac output was significantly higher in LNS8801-treated pregnant rats (67.3 ± 1.2 mL/min) compared to vehicle-treated pregnant rats (51.7 ± 4.6 mL/min; Figure 3C). Two-way ANOVA identified a significant main effect of treatment (*F*(1, 20) = 6.217, *p* = 0.0215, 21.35% of variation), while neither pregnancy (*F*(1, 20) = 0.6058, *p* = 0.4455, 2.08%) nor the interaction between treatment and pregnancy (*F*(1, 20) = 1.924, *p* = 0.1807, 6.61%) showed significant effects (Figure 3C). In virgin rats, no significant differences were observed in cardiac output (Figure 3C). Oral treatment of virgin and pregnant Dahl SS/Jr rats with the GPER agonist did not result in significant changes in cardiac structural indices, as shown in Table 1.

**Figure 3:**
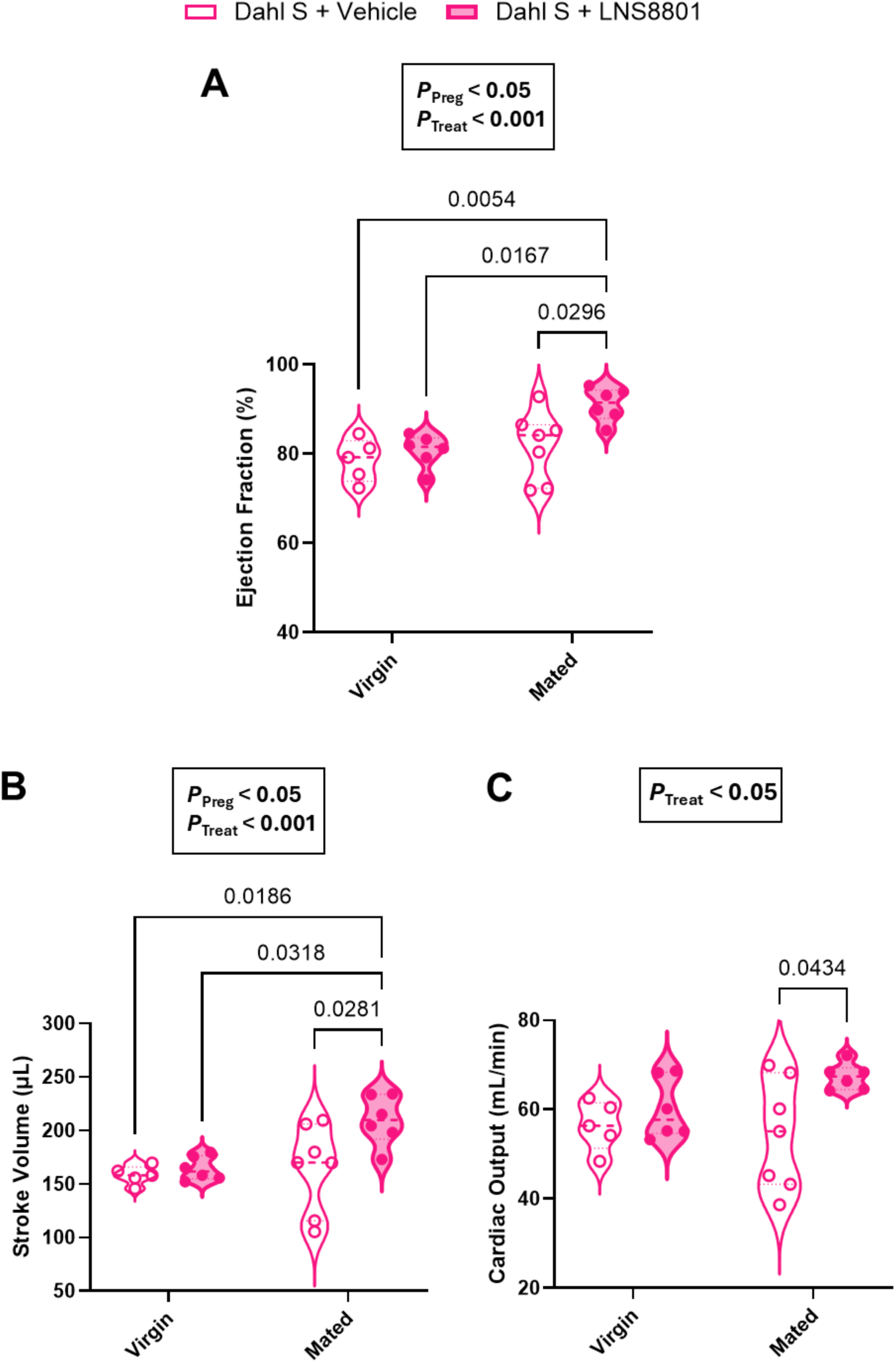
GPER activation with LNS8801 improves systolic function and cardiac output selectively in pregnant Dahl SS/Jr rats. (**A**) Ejection fraction was significantly increased in LNS8801-treated pregnant rats compared to all other groups, with no differences observed between virgin animals. (**B**) Stroke volume was higher in pregnant rats treated with LNS8801 compared to their respective controls and both virgin groups. (**C**) Cardiac output was also increased by LNS8801 in pregnant rats, while no effect was detected in virgin animals. Significant main effects of treatment and/or interaction (pregnancy × treatment) from two-way ANOVA are shown above each graph, and Tukey’s post hoc comparisons are indicated between groups, with significance set at (*p* < 0.05). Data are presented as mean ± SEM. Individual points represent biological replicates (n = 5 for Virgin + Vehicle, n = 6 for Virgin + LNS8801, n = 7 for Mated + Vehicle, and n = 6 for Mated + LNS8801).

### Effects of GPER stimulation with LNS8801 on blood pressure and proteinuria in virgin and pregnant Dahl SS/Jr rats

Our laboratory has shown that during late pregnancy, normal Sprague-Dawley rats had a terminal mean arterial pressure (MAP; measured under anesthesia) of 100.1 ± 3.6 mmHg^26^. In this study, terminal mean arterial pressure (MAP) was significantly reduced in pregnant Dahl SS/Jr rats treated with LNS8801 (122.9 ± 1.6 mmHg) compared to vehicle-treated pregnant controls (158.3 ± 3.0 mmHg), virgin rats treated with vehicle (169.9 ± 6.0 mmHg), and virgin rats treated with LNS8801 (167.6 ± 6.9 mmHg; Figure 4A). In virgin Dahl SS/Jr rats, LNS8801 treatment did not result in significant changes in MAP (Figure 4A). Two-way ANOVA for MAP revealed significant main effects of pregnancy (*F*(1, 20) = 35.56, *p* < 0.0001, 42.82%) and treatment (*F*(1, 20) = 16.01, *p* = 0.0007, 19.28%), as well as a significant interaction (*F*(1, 20) = 12.26, *p* = 0.0022, 14.77%). Additionally, urinary protein levels were significantly reduced in LNS8801- treated virgin rats (132.7 ± 7.9 mg/day) compared to virgin vehicle-treated controls (202.5 ± 26.4 mg/day), and in pregnant rats treated with LNS8801 (123.1 ± 7.9 mg/day) compared to their vehicle-treated counterparts (185.4 ± 16.6 mg/day; Figure 4B).

**Figure 4:**
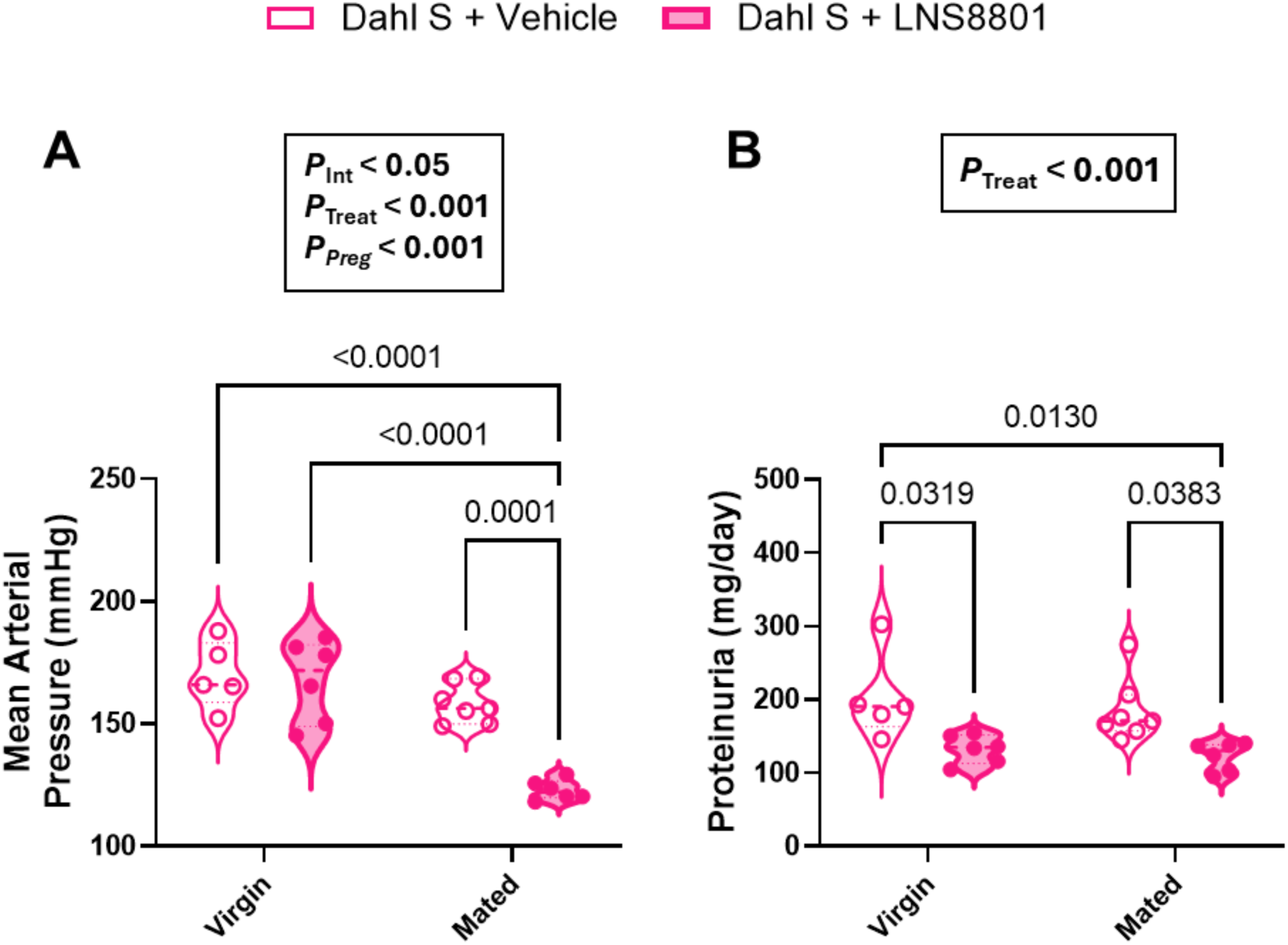
LNS8801 lowers mean arterial pressure in pregnant but not virgin Dahl SS/Jr rats, while reducing proteinuria in both. (**A**) Pregnant rats treated with LNS8801 showed a significant reduction in mean arterial pressure compared to all other groups. In virgin animals, LNS8801 had no significant effect on blood pressure. (**B**) LNS8801 treatment reduced urinary protein levels in both virgin and pregnant rats. The decrease was significant when compared to the respective vehicle-treated groups, and pregnant rats treated with LNS8801 also had lower proteinuria than virgin rats receiving vehicle. Significant main effects of treatment and/or interaction (pregnancy × treatment) from two-way ANOVA are shown above each graph, and Tukey’s post hoc comparisons are indicated between groups, with significance set at (*p* < 0.05). Data are presented as mean ± SEM. Individual points represent biological replicates (n = 5 for Virgin + Vehicle, n = 6 for Virgin + LNS8801, n = 7 for Mated + Vehicle, and n = 6 for Mated + LNS8801).

LNS8801-treated pregnant rats also exhibited significantly lower proteinuria than virgin vehicle-treated animals (Figure 4B). Two-way ANOVA confirmed a significant main effect of treatment (*F*(1, 20) = 17.65, *p* = 0.0004, 46.56% of the total variation), with no significant effects of pregnancy (*F*(1, 20) = 0.7188, *p* = 0.4066, 1.90%) or interaction (*F*(1, 20) = 0.05748, *p* = 0.8130, 0.15%).

### Impact of LNS8801 on fetal and placental outcomes

Previous studies have shown that pregnant Dahl SS/Jr rats tend to have smaller pups and higher rates of fetal resorptions compared to healthy pregnant rats^18^. In this work, treatment with LNS8801 did not impair fetal development, as pup weight remained consistent across the experimental groups (Figure 5A). Although LNS8801 did not significantly reduce fetal resorption rates (Figure 5B), it led to a significant reduction in placental weight in preeclamptic Dahl SS/Jr dams (0.39 ± 0.01 g) compared to the vehicle-treated group (0.53 ± 0.03 g; Figure 5C). This reduction in placental weight may have contributed to the significantly improved placental efficiency, which reflects the balance between fetal and placental weight, in LNS8801-treated rats (5.77 ± 0.38) compared to the control group (4.20 ± 0.53; Figure 5D).

**Figure 5:**
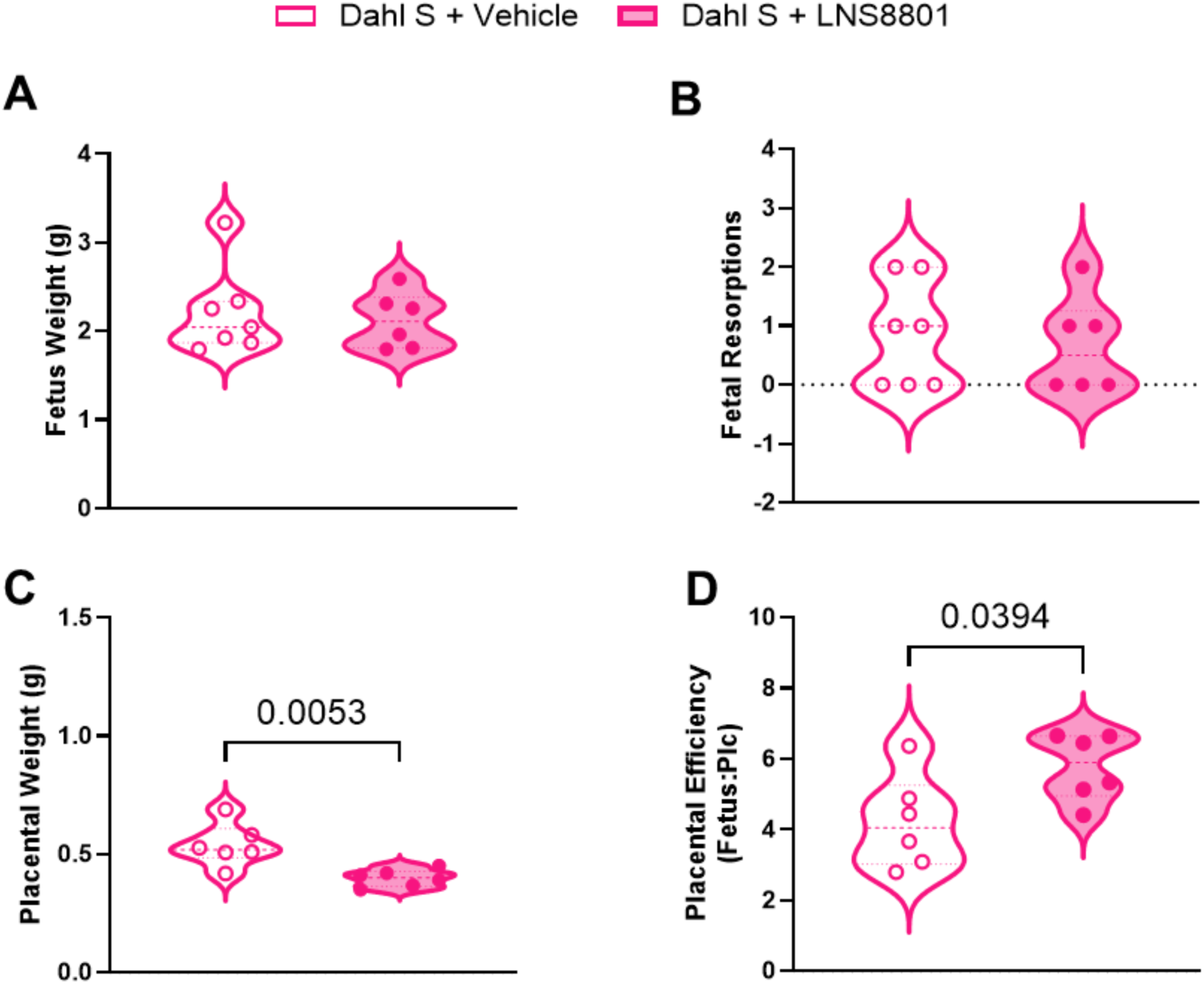
Fetal and placental outcomes following LNS8801 administration, assessed post-mortem on GD20. (**A**) Fetal weight was not significantly different between the LNS8801-treated and vehicle-treated groups. (**B**) The number of fetal resorptions was also not significantly different between the groups. (**C**) Placental weight was significantly lower in the LNS8801-treated group compared to the vehicle-treated group. (**D**) Placental efficiency, calculated as the ratio of fetal to placental weight (Fetus/Pc), was significantly higher in the LNS8801-treated group. Data is presented as mean ± SEM, with 6-7 experiments for the Dahl SS/Jr + vehicle group and 6 for the Dahl SS/Jr + LNS8801 group. Statistical analysis was conducted using an unpaired two-tailed t-test to compare differences between the experimental groups. A statistically significant difference between the Dahl SS/Jr + vehicle and Dahl SS/Jr + LNS8801 groups was considered at (*p* < 0.05).

## Discussion

This study demonstrates that activation of GPER with the selective agonist LNS8801 improves cardiovascular function in a pregnancy-dependent manner in hypertensive Dahl SS/Jr rats. In pregnant animals, LNS8801 enhanced both systolic and diastolic performance, normalized myocardial strain, and increased cardiac output - effects that were not observed in virgin rats - indicating that GPER-mediated cardioprotection requires the physiological context of pregnancy. These functional improvements were accompanied by reduced circulating levels of cardiac troponin I, consistent with decreased myocardial injury. LNS8801 also significantly lowered blood pressure and proteinuria in pregnant rats, addressing key features of superimposed preeclampsia. In virgin rats, while GPER activation with LNS8801 did not significantly impact cardiac function or blood pressure, it did reduce proteinuria, suggesting that the renal effects of GPER activation may be independent of pregnancy, whereas its cardiovascular benefits are pregnancy-specific.

LNS8801 is a highly selective GPER agonist developed by Linnaeus Therapeutics, containing only the active enantiomer responsible for GPER activation by G-1^22^. It is currently under investigation in clinical trials for its potential in cancer therapy (NCT04130516). Our investigation of LNS8801 effects in both pregnant and virgin Dahl SS/Jr rats highlights its therapeutic promise beyond cancer, particularly in GPER-related conditions such as pregnancy-associated cardiovascular dysfunction.

One of the central findings of this study is that activation of GPER by LNS8801 significantly improved myocardial function in pregnant, but not virgin, Dahl SS/Jr rats. Treatment significantly restored global longitudinal, circumferential, e radial strain toward physiological values, indicating enhanced myocardial deformation and mechanical efficiency in the hypertensive pregnant state. In parallel, systolic function was also amplified in pregnant rats receiving LNS8801, as shown by increases in ejection fraction and stroke volume, ultimately leading to greater cardiac output. These functional gains were accompanied by reductions in both end-diastolic and end-systolic volumes, reflecting more efficient cardiac filling and ejection dynamics. Together, these changes suggest enhanced cardiac performance likely supported by improved coordination between ventricular function and vascular load, consistent with GPER-mediated cardioprotection during pregnancy. In contrast, virgin animals showed no significant changes in strain or systolic parameters following LNS8801 administration, supporting the idea that the cardioprotective actions of GPER activation depend on the hemodynamic and hormonal milieu of pregnancy. Collectively, these findings indicate that LNS8801 enhances myocardial function specifically under conditions of pregnancy-associated cardiovascular stress, as seen in superimposed preeclampsia^27,28^.

The reduction in circulating cardiac troponin I levels exclusively in LNS8801-treated pregnant rats underscores the context-specific cardioprotection induced by GPER activation. Cardiac troponin I, a sensitive and specific biomarker for myocardial injury, is commonly used to detect cardiomyocyte stress and necrosis^29^. This selective effect suggests that the myocardial benefits of GPER stimulation become apparent only under the unique physiological load imposed by gestation. Although troponin levels remained unchanged in virgin animals, their decrease in pregnant rats implies that the therapeutic potential of LNS8801 may depend on the altered cardiac environment associated with late pregnancy complicated by hypertension. This is consistent with our myocardial strain data, further supporting the idea that GPER mitigates myocardial injury through improved mechanical performance when the heart is exposed to increased functional demand. The positive correlation between cardiac troponin I and global longitudinal strain strengthens this mechanistic link, indicating that reductions in myocardial stress are coupled with lower myocardial deformation. These findings align with prior research showing troponin I as a marker of myocardial stress in preeclamptic rats with cardiac dysfunction^30^.

In addition to its effects on myocardial strain and systolic function, LNS8801 also improved diastolic parameters in pregnant Dahl SS/Jr rats, as evidenced by increased early filling velocities (E wave) and a higher E/A ratio, indicating better ventricular relaxation. These changes were accompanied by a reduction in the E/e′ ratio, suggesting lower LV filling pressures. Impaired relaxation is a common feature of preeclamptic hearts^31^, and restoring this function would reduce the cardiovascular burden in affected pregnancies. In virgin Dahl SS/Jr rats, LNS8801 did not significantly alter E/A and E/e′ ratios, indicating that the beneficial impact of GPER activation on diastolic performance is more evident in the presence of pregnancy-induced cardiac stress.

Maternal hypertension is a hallmark of preeclampsia and often necessitates iatrogenic premature delivery^32^. In our study, the administration of LNS8801 at 800 µg/kg/day significantly reduced maternal blood pressure by 34 mmHg in pregnant Dahl SS/Jr animals, indicating its efficacy in managing hypertension in rats with superimposed preeclampsia. However, LNS8801 did not lower blood pressure in virgin Dahl SS/Jr rats, which aligns with previous studies showing that selective GPER stimulation does not reduce blood pressure in non-ovariectomized, non-pregnant rats with chronic hypertension^14–17^. This suggests that its antihypertensive effects are pregnancy-dependent, potentially influenced by pregnancy-specific vascular adaptations or hormonal interactions. Additionally, LNS8801 demonstrated significant renal benefits, as evidenced by a reduction in proteinuria in both pregnant and virgin Dahl SS/Jr rats. This reduction occurred independently of blood pressure changes in virgin animals, reinforcing that GPER activation directly influences renal function beyond its antihypertensive effects. This is consistent with previous findings showing that selective GPER stimulation with G-1 ameliorates kidney injury and reduces proteinuria in non-ovariectomized, non-pregnant Dahl SS/Jr rats without significantly altering blood pressure^17^.

The Dahl SS/Jr rat model, which is known to exhibit fetal growth restriction due to placental insufficiency^18^, mirrors the reduced fetal weights observed in preeclamptic pregnancies^33,34^. In our study, the fetal weight in the vehicle-treated group was also reduced, further supporting this model for studying the impacts of placental insufficiency. Interestingly, LNS8801 did not significantly alter fetal resorption rates or fetal weight, but it did reduce placental weight, which indicates improved placental efficiency. Enhancing placental efficiency is essential in the context of preeclampsia, where placental abnormalities play a major role in adverse outcomes^35^.

Our previous research demonstrated the cardioprotective effects of GPER activation using G-1 in the reduced uterine perfusion pressure model of preeclampsia^20^, which has been widely used to study preeclampsia-related cardiovascular dysfunction. However, the Dahl SS/Jr rat model of superimposed preeclampsia used in this study provided additional insights into how chronic hypertension interacts with pregnancy-related vascular and cardiac adaptations. The presence of chronic hypertension is particularly relevant, as many individuals who develop preeclampsia already have underlying cardiovascular risk factors. By investigating both virgin and pregnant Dahl SS/Jr rats, this study also allows us to explore how pregnancy modulates GPER activation in a hypertensive setting. Furthermore, LNS8801 presents a clinically viable alternative to G- 1, offering the advantages of oral bioavailability and ongoing clinical trials, which provide an established safety profile that could accelerate its translation into preeclampsia treatment.

In conclusion, LNS8801 demonstrates substantial benefits in improving cardiovascular and renal function in a preclinical model of chronic hypertension in pregnancy, with distinct effects that depend on pregnancy status. Its pharmacological advantages, such as oral bioavailability and high selectivity for GPER, support its potential for further development as a non-invasive therapeutic option for preeclampsia. These findings open new avenues for targeted treatments that address both cardiovascular and renal dysfunction, which remain critical challenges in the management of high-risk pregnancies.

## Limitations of the Study

Although previous studies have reported further elevations in blood pressure during pregnancy in Dahl SS/Jr rats, such findings were primarily based on longitudinal assessments comparing pre-pregnancy and late-gestation time points within the same animals. In contrast, our study employed a cross-sectional design comparing age-matched virgin and pregnant animals. As a result, we did not observe significantly higher mean arterial pressure in pregnant animals compared to their virgin counterparts. Nevertheless, all animals in our study were hypertensive, as expected for this strain, and pregnant animals exhibited exacerbated cardiac dysfunction, including altered systolic and diastolic performance and increased myocardial strain, compared to virgins. These results support the use of this model to investigate superimposed heart stress during pregnancy, even in the absence of additional elevations in arterial pressure.

While this study demonstrated clear improvements in cardiac and renal function following GPER activation, it did not explore the underlying molecular mechanisms - such as inflammation or oxidative stress - that may contribute to these functional changes. Future studies will be necessary to define the mechanistic pathways involved and expand upon the physiological findings reported here. Finally, although LNS8801 is currently under clinical investigation for oncology indications, its safety, efficacy, and pharmacokinetics during pregnancy remain unknown and must be thoroughly evaluated before translational application in the context of preeclampsia.

## Perspectives and Significance

GPER activation during pregnancy plays a crucial role in modulating cardiovascular and renal adaptations, which may have therapeutic implications for hypertensive disorders of pregnancy. In this study, we investigated the effects of LNS8801, a selective and orally bioavailable GPER agonist, in virgin and pregnant Dahl SS/Jr rats, a model of chronic hypertension that progresses to pregnancy-related cardiovascular dysfunction.

Our findings reveal that pregnancy-specific adaptations enhance GPER-mediated cardioprotection, as LNS8801 significantly improved cardiac function in pregnant rats, whereas its effects were limited in virgin animals.

Additionally, LNS8801 reduced proteinuria in both groups, independent of blood pressure changes in virgins, suggesting a broader renal protective role for GPER activation beyond pregnancy. These results highlight the potential of GPER-targeted therapies for managing preeclampsia, particularly in populations with pre-existing hypertension. Given that LNS8801 is already in clinical trials for cancer, our study positions this compound as a promising candidate for future preeclampsia treatments. Furthermore, this research paves the way for exploring genetic factors and long-term cardiovascular risks for both mothers and offspring, advancing both preventative and therapeutic strategies for this high-risk population.

## Novelty and Relevance

### What Is New?

- GPER (G protein-coupled estrogen receptor) is a novel therapeutic target for managing cardiovascular and renal complications in superimposed preeclampsia, with its effects shaped by pregnancy-specific physiological adaptations.
- Activation of GPER by the selective agonist LNS8801 significantly improves cardiac function and reduces blood pressure in pregnant Dahl SS/Jr rats with superimposed preeclampsia, whereas its effects on cardiac function and blood pressure were limited in virgin animals.
- LNS8801 reduces proteinuria in both virgin and pregnant Dahl SS/Jr rats, suggesting a pregnancy-independent role for GPER activation in renal protection, which could extend its therapeutic potential beyond preeclampsia.
- LNS8801, currently undergoing clinical trials for cancer, has already established a safety profile in humans, making it a strong candidate for repurposing in preeclampsia treatment.
- The Dahl SS/Jr rat offers a clinically relevant model of superimposed preeclampsia, capturing both chronic hypertension and pregnancy-induced cardiorenal dysfunction.

### What Is Relevant?

GPER activation by LNS8801 reveals that pregnancy-specific adaptations influence cardiovascular and renal function in a preclinical model of superimposed preeclampsia. While LNS8801 improved cardiac performance and reduced hypertension in pregnant Dahl SS/Jr rats, these benefits were not observed in virgin animals. In contrast, LNS8801 reduced proteinuria in both groups, suggesting a broader renal protective effect that is independent of pregnancy status and may be relevant to hypertensive disorders beyond preeclampsia. Using the Dahl SS/Jr rat model, which naturally develops hypertension before pregnancy, this study provides a clinically relevant framework to investigate how pre-existing cardiovascular dysfunction interacts with pregnancy-specific adaptations in the context of preeclampsia and potential therapeutic interventions. The differential responses observed between virgin and pregnant animals indicate the need for pregnancy-specific models in translational research. These findings contribute to a more nuanced understanding of GPER activation as a therapeutic strategy, with potential implications for managing both pregnancy-associated and chronic hypertensive disorders.

## Acknowledgements

Not applicable.

## Author contributions

C.L.B. and A.K.N.A. conceived and designed the research. A.K.N.A., K.F.S., and M.M.M. performed experiments and analyzed data. C.L.B., A.K.N.A., and K.F.S. interpreted the results of experiments, prepared figures, and drafted the manuscript. C.L.B., A.K.N.A., K.F.S., S.H.L., C.A.N., G.C.P., and M.R.G. edited and revised the manuscript and approved the final version.

