## Supplemental Material for "Pregnancy-Dependent Cardioprotection via GPER Activation in Dahl Salt-Sensitive Rats"

**Short title:** Pregnancy-Specific Cardiac Effects of GPER Activation in Rats

1. Department of Biomedical Engineering, Tulane University, New Orleans, LA, 70118, USA.

2. Department of Obstetrics & Gynecology, Tulane University, New Orleans, LA, 70112, USA.

3. Linnaeus Therapeutics, Haddonfield, NJ, USA.

4. Department of Pharmacology, Tulane University, New Orleans, LA, 70112, USA.

5. Department of Cell and Molecular Biology, University of Mississippi Medical Center, Jackson, MS, USA.

***Correspondence and reprint requests to:** Carolyn L. Bayer, Department of Biomedical Engineering, Tulane University, 500 Lindy Boggs Center, New Orleans, LA, 70118, USA..

### Material and Methods

#### *Biochemical Analyses*

The Cardiac Troponin I circulating levels were measured in this study. Blood samples were collected on GD20 using sterile centrifuge tubes immediately prior to euthanasia via the catheter inserted into the left common carotid artery. The collected blood was allowed to coagulate, followed by centrifugation, and the resulting serum was aliquoted. These serum aliquots were then stored at -80 °C until further processing. The measurement of Cardiac Troponin I levels was conducted using an enzyme-linked immunosorbent assay (ELISA) in strict accordance with the instructions provided by the manufacturer (R&D Systems, Minneapolis, MN).

### Supplemental Figures

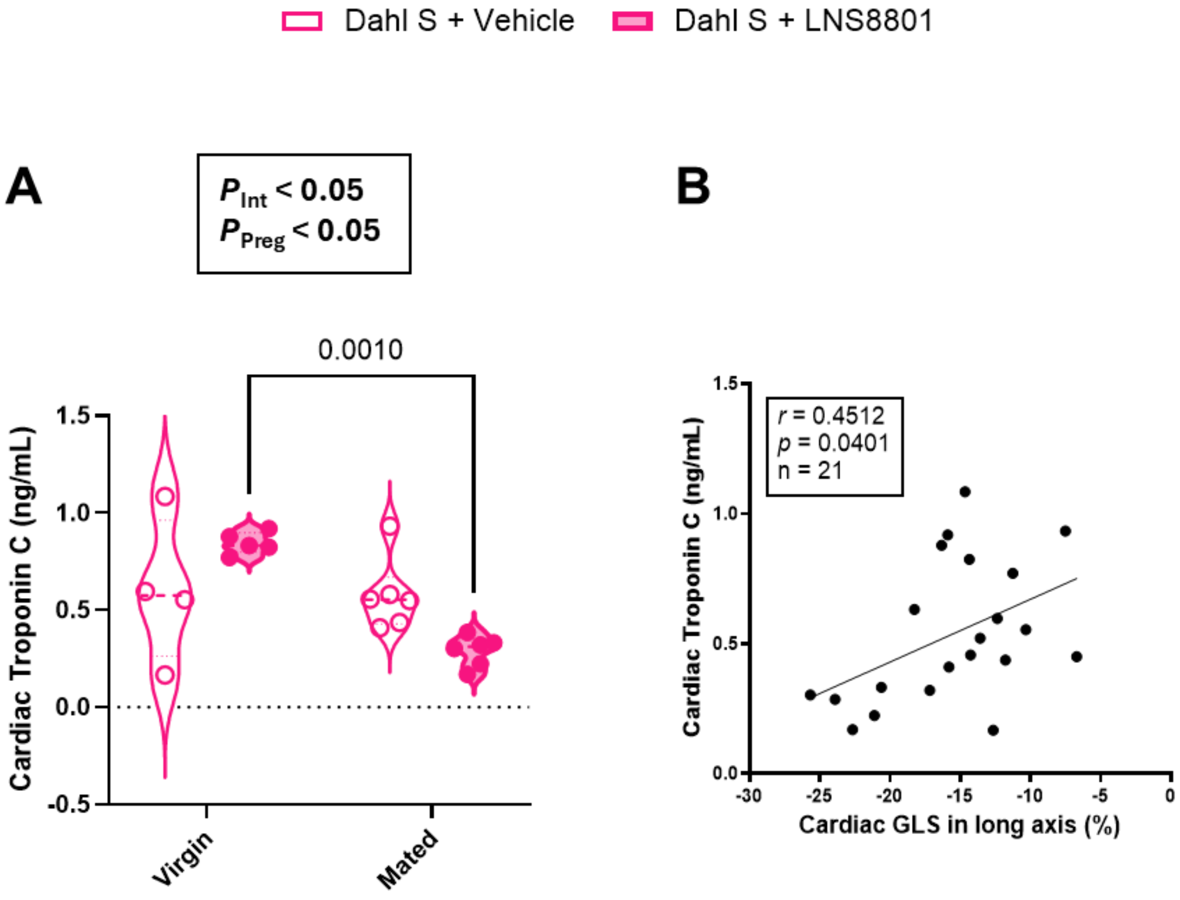

**Figure S1:** GPER activation by LNS8801 normalized circulating levels of cardiac troponin I in pregnant but not virgin Dahl SS/Jr rats. (**A**) Circulating levels of cardiac troponin I were significantly higher in mated animals treated with LNS8801 (800 µg/kg/day) compared to the mated group treated with the vehicle. No significant difference was observed between treatment groups in virgin animals. (**B**) Cardiac troponin I levels were positively correlated with global longitudinal strain (GLS) in the long axis, suggesting that increased troponin release is associated with cardiac dysfunction in this model. Data is presented as mean ± SEM. Due to three hemolyzed samples, the final sample sizes for this dataset were Virgin + Vehicle (n = 4), Virgin + LNS8801 (n = 5), Pregnant + Vehicle (n = 6), and Pregnant + LNS8801 (n = 6), slightly reduced compared to other *in vivo* datasets. Statistical analysis was performed using two-way ANOVA followed by Tukey's multiple comparisons test. Significant main effects of pregnancy and interaction (pregnancy × treatment) from two-way ANOVA are shown above the graph, and Tukey's post hoc comparisons are indicated between groups.
